# Functional Modulation of Retrotrapezoid Neurons Drives Fentanyl-Induced Respiratory Depression

**DOI:** 10.1101/2025.01.28.635295

**Authors:** Thiago S. Moreira, Nicholas J. Burgraff, Ana C. Takakura, Luiz M. Oliveira, Emmanuel V. Araujo, Steven Guan, Jan-Marino Ramirez

**Affiliations:** Department of Physiology and Biophysics, Institute of Biomedical Science, University of Sao Paulo, Sao Paulo, Brazil; Center for Integrative Brain Research, Seattle Children’s Research Institute, Seattle, WA, USA; Department of Comparative Biosciences, School of Veterinary Medicine, University of Wisconsin-Madison, Madison, WI, USA; Department of Pharmacology, Institute of Biomedical Science, University of Sao Paulo, Sao Paulo, Brazil; Department of Neurological Surgery, University of Washington, Seattle, WA, USA

**Keywords:** parafacial region, breathing, chemoreceptors, opioids

## Abstract

The primary cause of death from opioid overdose is opioid-induced respiratory depression (OIRD), characterized by severe suppression of respiratory rate, destabilized breathing patterns, hypercapnia, and heightened risk of apnea. The retrotrapezoid nucleus (RTN), a critical chemosensitive brainstem region in the rostral ventrolateral medullary reticular formation contains Phox2b^+^/Neuromedin-B (*Nmb*) propriobulbar neurons. These neurons, stimulated by CO_2_/H^+^, regulate breathing to prevent respiratory acidosis. Since the RTN shows limited expression of opioid-receptors, we expected that opioid-induced hypoventilation should activate these neurons to restore ventilation and stabilize arterial blood gases. However, the ability of the RTN to stimulate ventilation during OIRD has never been tested. We used optogenetic and pharmacogenetic approaches, to activate and inhibit RTN Phox2B^+^/*Nmb*^+^ neurons before and after fentanyl administration. As expected, fentanyl (500 µg/kg, ip) suppressed respiratory rate and destabilized breathing. Before fentanyl, optogenetic stimulation of Phox2b^+^/*Nmb*^+^ or chemogenetic inhibition of *Nmb*^+^ cells increased and decreased breathing activity, respectively. Surprisingly, optogenetic stimulation after fentanyl administration caused a significantly greater increase in breathing activity compared to pre-fentanyl levels. By contrast chemogenetic ablation of RTN *Nmb* neurons caused profound hypoventilation and breathing instability after fentanyl. The results suggest that fentanyl does not inhibit the ability of Phox2b^+^/*Nmb*^+^ cells within the RTN region to stimulate breathing. Thus, this study highlights the potential of stimulating RTN neurons as a therapeutic approach to restore respiratory function in cases of OIRD.

## Introduction

The escalating crisis of opioid overdoses highlights a critical medical challenge, i.e opioid-induced respiratory depression (OIRD), the leading cause of mortality in such cases (1, 2). OIRD tragically disrupts normal respiratory functions, leading to decreased respiratory rate, altered breathing patterns, severe hypercapnia, and an increased risk of fatal apnea (1, 3). Fentanyl, a synthetic opioid, is notably more potent than morphine, as highlighted in several studies (4, 5). Growing concerns suggest that fentanyl may present higher risks for OIRD compared to other opioid treatments, especially due to its tendency to cause chest wall rigidity and laryngeal closure - a phenomenon known as ‘wooden chest syndrome,’ observed in both humans and rodents (6–9). This condition is less commonly linked with morphine and other opioid analgesics, highlighting the need for a deeper understanding of the distinct impacts of these opioids (10).

The respiratory network receives excitatory chemosensitive drive from the carotid body (CB), via the nucleus of the solitary tract (NTS), and from the retrotrapezoid nucleus (RTN) and from many other nuclei such as medullary raphe, locus coeruleus and lateral hypothalamus (11–15). The carotid body acts as a peripheral chemoreceptor, sensing changes in blood oxygen (O_2_) levels and communicating this information to the brainstem via the NTS, which adjusts eupneic breathing patterns (16). Hypoxia also increases the sigh rate (17–20), and the NTS also plays a pivotal role in integrating and processing respiratory and cardiovascular signals (21, 22). While the CB is particularly sensitive to hypoxia, the RTN, located within the rostral ventrolateral medullary reticular formation (13, 23), plays a pivotal role in controlling respiratory responses to elevated levels of carbon dioxide (CO_2_/H^+^) (24–26). The RTN is key to maintaining normal breathing patterns under various physiological conditions (27–29). Therefore, all these regions (CB, NTS, and RTN) contain specialized neurons that express the paired-like homeobox 2B transcription factor (Phox2b). However, the RTN exhibits limited expression of opioid receptors, highlighting its distinct role and resilience to the direct effects of opioids (30). These neurons, inherently sensitive to CO_2_/H^+^ environments, are fundamental in stimulating respiratory activity and thereby averting respiratory acidosis under typical circumstances such as the OIRD (24, 25, 31, 32). Importantly, the activation of these neurons by elevated CO_2_/H^+^ functions as a critical feedback mechanism that should, in theory, counteract opioid-induced respiratory depression by enhancing ventilation to normalize arterial blood gases.

However, the efficiency and mechanism of all these chemosensitive neurons in counteracting the depressive respiratory effects during an opioid overdose remain inadequately understood. This gap in knowledge presents a significant barrier to effectively addressing OIRD. Their ability to monitor hypoxia and hypercapnia could serve as a potential lifeline, restoring normal respiratory function, eupneic breathing and sighing amidst the life-threatening conditions induced by opioid toxicity. In this study, we aimed to investigate more deeply the role of chemosensitive neurons in the depression of normal breathing (eupnea) and the generation of sighs under the influence of fentanyl. In addition, it is well described that central chemoreceptor neurons within the RTN region can be identified by the expression of a constellation of markers, including expression of Phox2b, vesicular glutamate transporter 2 (Slc17a6), tachykinin receptor 1 (TACR1), galanin, and more recently neuromedin B (*Nmb*), as well as the absence of TH and ChAT (31). Among these identifiers, *Nmb* is universally expressed in putative central chemoreceptor neurons of the RTN (17, 33). Thus, we also hypothesized that *Nmb* neurons in the RTN contribute to respiratory homeostasis and are essential for rescuing breathing during OIRD.

## Material and methods

### Animals

All experimental and surgical procedures were conducted in accordance with the guidelines set by the National Institutes of Health guidelines (NIH Publications No. 8023, revised 1978), Seattle Children’s Research Institute Animal Care and Use Committee (protocol IACUC00058), and Animal Care and Use Committees of the University of São Paulo (Protocol # CEUA-ICB/USP: n° 9750170720). Male (N = 20) and female (N = 21) mice weighing between 20-28g were used. The animals were provided with *ad libitum* access to water and food and were housed under controlled environmental conditions, including a temperature of 23 ± 1 °C, humidity of 50 ± 10%, and a 12-hour light-dark cycle (lights on at 07:00 a.m).

We used transgenic Phox2b::cre mice (*B6(Cg)-Tg(Phox2b-cre)3Jke/J* – JAX#016223) crossed with homozygous Ai32 mice (Jax# 012569) that contain a flox-STOP-flox sequence fused to channelrhodopsin (ChR2) and enhanced yellow fluorescent protein at the Rosa26 locus (Figs. 1A-B). Thus, offspring exhibit ChR2 expression and enhanced yellow fluorescent protein only in the neurons that express the promoter for the cre driver. We also used transgenic *Nmb*::cre^/+^ mice. The *Nmb*::cre mouse line was maintained in house by crossing with C57BL/6J mice acquired from The Jackson Laboratory (stock #000664), and male and female WT control (*Nmb*^+/+^) and heterozygous cre-positive littermates (*Nmb*::cre^/+^) were used for experiments.

**Figure 1).**
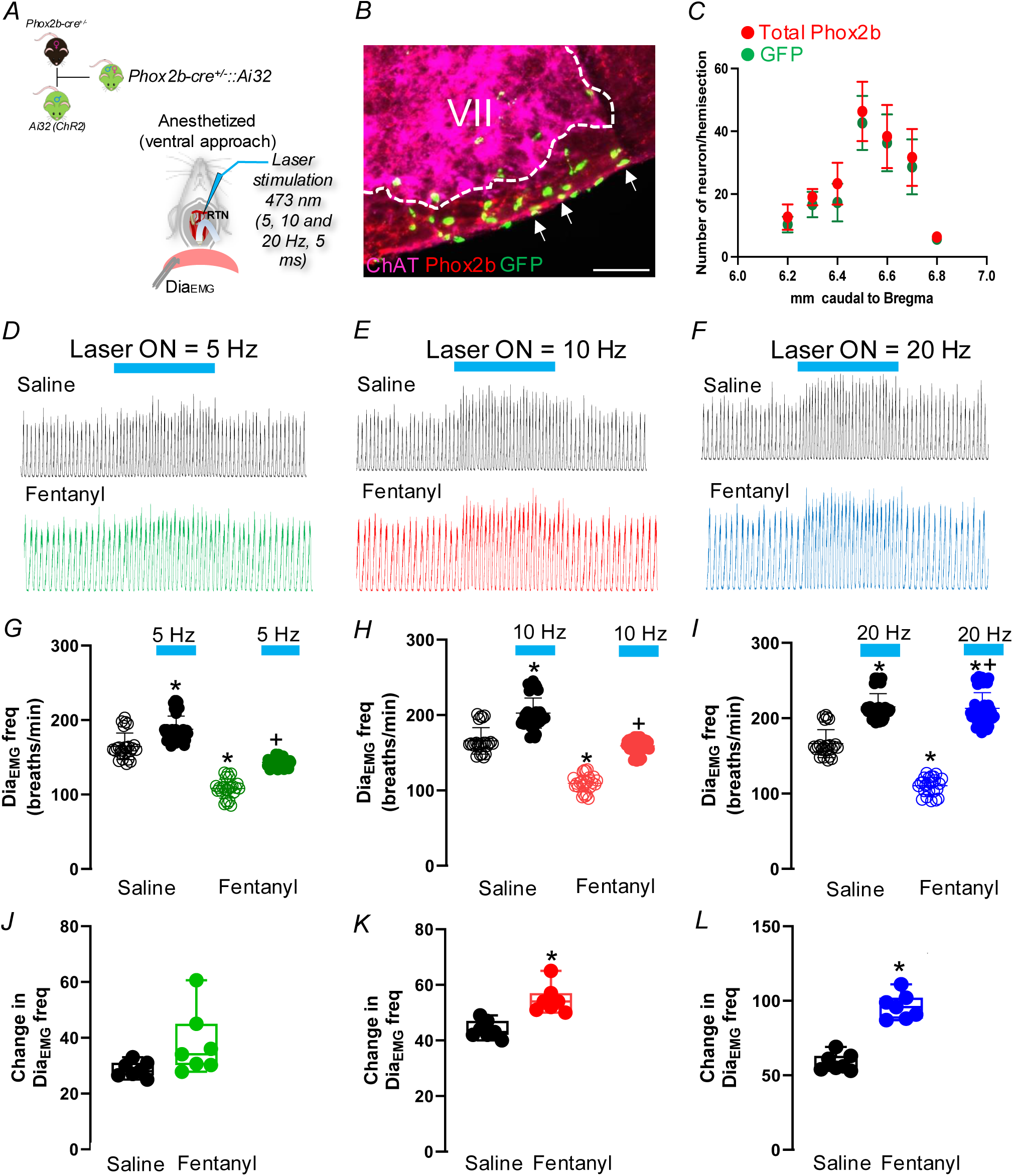
Optogenetic activation of ChR2-expressing Phox2b RTN neurons activates breathing in anesthetized mice treated with fentanyl. A) Diagram showing how we generate a conditional Phox2b::cre^+/-^;Ai32. Schematic of the *in vivo* (urethane anesthetized) preparation. In anesthetized mice, diaphragm muscle was recorded and a ventral optrode was placed in the RTN region. B) Immunohistochemistry showing the co-localization of Phox2b with GFP fluorescence. C) Total number of Phox2b^+^ neurons transduced in the RTN region. D-F) Representative recordings of diaphragm electromyography (Dia_EMG_) from an anesthetized mouse after ip injection of saline or fentanyl (500 μg/kg) under baseline or photostimulation (5, 10 and 20 Hz, 10 ms light pulse) of the Phox2b-expressing neurons in the RTN region. G-I) Group data for the Dia_EMG_ respiratory frequency effect of G) 5Hz (One way repeated-measure ANOVA F_3, 146_ = 200; p<0.0001), H) 10 Hz (One way repeated-measure ANOVA F_3, 146_ = 208; p<0.0001), and I) 20 Hz photostimulation of Phox2b-expressing RTN neurons in a saline or fentanyl injection (One way repeated-measure ANOVA F_3, 146_ = 236.5; p<0.0001). J-L) Changes in Dia_EMG_ respiratory frequency elicited by J) 5Hz, K) 10 Hz, and L) 20 Hz photostimulation of Phox2b-expressing RTN neurons in a saline or fentanyl injection. Abbreviation: VII, facial motor nucleus. Scale bar = 100 μm, applied to B. *Different from baseline (saline); +Different from baseline (fentanyl).

### Viral vectors

For optogenetic stimulation of RTN *Nmb* neurons, we used pAAV5-EF1a-double floxed-hChR2(H134R)-mCherry-WPRE-HGHpA (Addgene plasmid 20297, titer for injection: 1.0 × 10^13^ vg/ml). For chemogenetic inhibition, we used a cre-dependent vector containing the fluorophore mCherry [AAV5-hSyn-DIO-hM4D(Gi)-mCherry, Addgene plasmid 44362, titer: 7.0 × 10^12^ vg/ml].

### *In vivo* anesthetized preparation and physiological recordings

All surgical procedures and experimental protocols were done in urethane (1.2 g/kg, intraperitoneal - ip) anesthetized mice prepared as described previously (34–37). The anesthesia level was monitored by testing for the absence of withdrawal response and lack of diaphragm electromyography activity (Dia_EMG_) changes due to firm paw pinch.

The mice were placed supine on a custom surgical table. Body temperature was kept close to 37°C with a water heating system (PolyScience, Niles, IL) built into the surgical table. Mice were then allowed to spontaneously breathe 100% O_2_ or room air for the remainder of the surgery and experimental protocol.

The adaptor was outfitted with a nose mask (Kopf model 907) through which 100% O_2_ or room air was delivered. Dia_EMG_ was recorded by placing bipolar electromiography electrodes in the costal diaphragm to monitor respiratory rate throughout the experiment (34, 38). Signal processing occasionally included additional digital bandpass narrowing to minimize the electrocardiography artifact. This was followed by signal rectification and integration (0.03 s time constant). Respiratory frequency (f_R_) was triggered from the integrated Dia_EMG_ signal.

The trachea was exposed through a midline incision and cannulated caudal to the larynx with a curved (180°) tracheal tube (PTFE 24G, Component Supply, Sparta, TN). Continuing in the supine position, the occipital bone was removed, followed by continuous perfusion of the ventral medullary surface with warmed (∼36°C) artificial cerebral spinal fluid (aCSF; in mM: 118 NaCl, 3 KCl, 25 NaHCO3, 1 NaH2PO4, 1 MgCl2, 1.5 CaCl2, 30 d-glucose) equilibrated with carbogen (95% O_2_, 5% CO_2_) by a peristaltic pump (Dynamax RP-1, Rainin Instrument Co, Emeryville, CA). A fiber optic cable (200 μm diameter) connected to a blue (447 nm) laser and DPSS driver (Opto Engine LLC, Salt Lake City, UT) was placed unilaterally in light contact with the ventral surface of the brainstem overtop of the predetermined RTN region or in the carotid artery bifurcation where the carotid body was located (12, 35, 39, 40) (Figs. 1A and 3D - anesthetized ventral approach). All the experimental protocols were executed before and after the injection of fentanyl. Fentanyl was administered at 500 μg/kg (ip) from a stock solution of 50 μg/ml (Hikma Pharmaceuticals, London, UK). Doses of fentanyl were determined based upon the amount necessary to elicit suppression of respiratory frequency as previously demonstrated (9). The entire experiment, including surgery and recordings, lasted <3 h.

### *In vivo* non-anesthetized preparation and physiological recordings

General anesthesia was induced with 4% isoflurane in 100% O_2_. The depth of anesthesia was assessed by an absence of the hind-paw withdrawal reflex, then the anesthesia with 2-2.5% isoflurane in 100% O_2_ was maintained throughout surgery. The following procedures were performed under aseptic conditions. Following skin shaving and disinfection, the mice were then placed on a stereotaxic apparatus adapted for mouse. Body temperature maintained at 37°C with a servo-controlled heating pad and a blanket. A small hole, 1.5 mm in diameter, was drilled unilaterally into the occipital plate caudal to the parieto-occipital suture on the left side to implant the optical fiber vertically (0.2 mm core, 0.37 numerical aperture; Thorlabs). To photostimulate the Phox2b RTN cells the tip of the fiber optic was inserted 200-250 μm dorsal to where the Phox2b-expressing cells have been located (5.35 mm below the dorsal surface of the brain, 1.3 mm lateral to the midline, and 1.4 mm caudal to lambda). To photostimulate the Phox2b NTS cells the tip of the fiber optic was inserted 100-150 μm dorsal to where the Phox2b-expressing cells have been located (4 mm below the dorsal surface of the brain, 0.3 mm lateral to the midline, and 1.8 mm caudal to lambda).

For the optogenetic experiments in *Nmb*::cre^/+^ mice, a vector encoding channelrodopsin-ChR2 (AAV5-EF1a-double floxed-hChR2(H134R)-mCherry-WPRE-HGHpA) was microinjected in the left RTN in two sites spanning the rostro-caudal length of the facial nucleus (1.4 mm lateral to midline, 100 µm below the facial motor nucleus). After the vector injection, a fiber optic was implanted in the direction to the RTN region as described above. For the inhibition of RTN *Nmb* neurons, microinjections of AAV5-hSyn-DIO-hM4D(Gi)-mCherry, 60-80 nl of virus were delivered bilaterally at two sites spanning the rostro-caudal length of the facial nucleus (1.4 mm lateral, 100 µm below the facial motor nucleus).

Three-four weeks after the implantation of the guide cannula or vector injections, the mice were placed in the plethysmography chamber and habituated to chamber conditions for 2 h. At this point, mice were taken to allow for insertion of a custom optical fiber assembly into the implanted guide or injection of saline or the DREADD ligand CNO at 0.1 mg/kg i.p. Mice were returned to the plethysmography chamber and allowed to recover for an additional 30-40 min before commencing experimental protocols.

Breathing parameters were evaluated in conscious unrestrained mice using whole-body plethysmography (500 ml internal volume). Recording sessions were conducted between 8:00 am and 5:00 pm. Chambers were continuously flushed with dry room temperature air delivered by flow regulators controlling the flow of gases (O_2_ and CO_2_) (total flow: 0.5-1 L/min). The flow signal was amplified (×500) and acquired at 1 kHz in Spike2 software (version 7, CED). Chamber temperature, relative humidity, and atmospheric pressure were continuously monitored and were very stable within experiments (temperature; 24.5 ± 0.1°C, relative humidity; 42.4 ± 1.8%). Body temperature was measured before and after the experimental protocol and an average was calculated.

These mice were used to evaluate the effects of photostimulation or chemogenetic inhibition on breathing during normoxia (FiO_2_ = 0.21, balanced by N_2_), and 10 min challenges of hypercapnia (FiCO_2_ = 0.1) or hypoxia (FiO_2_ = 0.1, balanced by N_2_). For the normoxic hypercapnia protocol, the inspired fraction of oxygen (FiO_2_) was set at 0.21 and then stayed at this level during the whole protocol, similar to our previous studies (36, 37). After 20 min of normoxia, the inspired fraction of CO_2_ (FiCO_2_) increased from 0 to 0.1. The episode of hypercapnia lasted 10 min and was followed by recovery in normoxia for at least 20 min. The hypoxia protocol was performed by exposing the mice to FiO_2_ = 0.1 for 10 min, following a period of normoxia recovery (reoxygenation) for 20 min. Respiratory frequency (f_R_) was analyzed from periods free of movement artifacts. During hypercapnia and hypoxia, the analysis of f_R_ was performed at 7-10 min from the beginning of the gas change. The experimental protocol was executed before and after the injection of fentanyl (500 μg/kg, ip) (9).

At the completion of the experiment, these mice were anesthetized and perfused according to the methods detailed below.

### Histology analysis and cell counts

At the end of experiments, mice were deeply anesthetized with ketamine/xylazine (100/10 mg/kg, i.p.), after confirming the absence of the toe pinch reflex, 50 units of heparin were injected transcardially. The mice were then perfused through the ascending aorta with 50 ml of phosphate buffered saline (PB; pH 7.4) followed by 100 ml of 4% phosphate-buffered formaldehyde (0.1 M, pH 7.4) (Electron Microscopy Sciences). The brains were then removed and stored in fixative for 4 hours at 4°C and sucrose 20% overnight. A series of coronal sections (1:4 series, 30 μm thick) were cut along the rostrocaudal axis using a microtome (SM2010R; Leica Biosystems, Buffalo Grove, IL USA) and stored in cryoprotectant solution at −20°C (20% glycerol, 30% ethylene glycol in 50 ml PB) for later histological processing. All histochemical procedures were performed using free-floating sections, in accordance with previously described protocols (36, 37).

By immunofluorescence technique, tyrosine hydroxylase (TH) was detected using a polyclonal mouse anti-TH antibody (MAB 318; Millipore; 1:1000); Phox2b was detected using a polyclonal rabbit anti-Phox2b antibody (AB 2813765, Santa Cruz Biotechnology; 1:800); and choline acetyltransferase (ChAT) was detected using a polyclonal goat anti-ChAT antibody (AB 144P; Millipore; 1:100) diluted in PBS containing normal donkey serum (017-000-121; 1%; Jackson Immuno Research Laboratories) and Triton-X 0.3% and incubated for 24 h. The sections were subsequently washed in PBS and incubated for 2 hours in the donkey anti-mouse Alexa 594 (715-585-151; Jackson Immuno Research Laboratories; 1:500); donkey anti-rabbit Alexa 594 (711-585-152; Jackson Immuno Research Laboratories; 1-500); donkey anti-rabbit Alexa 488 (711-605-152; Jackson Immuno Research Laboratories; 1-500), and donkey anti-goat Alexa 647 (715-605-151; Jackson Immuno Research Laboratories; 1-500) were used when appropriated. Brain sections have been blinded analyzed using a fluorescence microscope (ZeissAxioskop A1, Oberkochen, Germany) for counting Phox2b neurons in RTN region.

We employed the RNAscope™ Multiplex Fluorescent Reagent Kit (Advanced Cell Diagnostics) for our analysis. Initially, brain sections were immersed in 0.1M PBS for 5 minutes, followed by a 10-minute treatment with RNAscope Hydrogen Peroxide (#322381) at room temperature (RT). Subsequently, the sections were rinsed three times with PBS, each rinse lasting 2 minutes. The tissue was then exposed to Protease IV (#322381) for 30 minutes at RT, followed by another 2-minute incubation in 0.1M PBS. Next, the sections were incubated with Slc17a6 and Nmb probes for 2 hours at 40°C, followed by a brief 2-minute PBS rinse. For hybridization, the tissue underwent a series of incubations with RNAscope Multiplex Detection Reagents (#323110), starting with FL v2 Amp 1 for 30 minutes at 40°C, with a 5-minute PBS wash after each step. Following this, the tissue was incubated with RNAscope Multiplex FL for 30 minutes at 40°C, followed by another 5-minute PBS rinse. The same was applied to FL v2 Amp 2. To visualize the HRP-C1 and HRP-C2 signals, the sections were treated with RNAscope Multiplex FL v2 HRP-C1 or HRP-C2 for 15 minutes at 40°C, followed by a 5-minute PBS wash. Signal amplification was achieved using TSA Plus fluorescein (FP1168; 1:250) for 30 minutes at 40°C, followed by another 5-minute PBS rinse. The tissue was then incubated with RNAscope Multiplex FL v2 HRP blocker for 15 minutes at 40°C, followed by a final 2-minute PBS wash. Finally, all slides were mounted using Fluoromount (00-4958-02; Thermo Fisher), and coverslips were secured with nail polish.

Brain sections have been blinded analyzed using StereoInvestigator software (MBF Bioscience) with a fluorescence microscope (ZeissAxioskop A1, Oberkochen, Germany). Images were acquired with a Hamamatsu C11440 Orca-Flash 4.0LT digital camera (resolution: 2048 x 2048 pixels) resulting in TIFF files. A technical illustration software package (Adobe Illustrator, v. 28.1, USA) was used for line drawings, figure assembly, and labeling, following the guidelines of Franklin & Paxinos (41). Representative images were pseudo-colored and optimized for presentation, brightness, and contrast were adjusted equally in all pixels of the image. Images were manually cropped to the relevant anatomic region, a Gaussian blur was applied, the default auto thresholding was applied (42) despeckled, and then the default particle analysis tool was used to collect signal intensities. The files were exported into the Canvas drawing program (version 9, Deneba Systems Inc., Miami, FL) for text labeling and final presentation.

The sections were counted bilaterally, and the numbers reported in the results section correspond exactly to the counts of one-in-four sections in a series. Section alignment between brains was completed relative to a reference section, as previously described (33, 37, 43). Briefly, to align sections around the RTN level, the most caudal section that contained an identifiable cluster of facial motor neurons was identified in each brain and assigned to the level of 6.48 mm caudal to the bregma (bregma = - 6.48 mm). To align sections around the NTS level, the most caudal section was identified in each brain and assigned to the level 7.92 mm caudal to the bregma (bregma = - 7.92 mm). Levels rostral or caudal to this reference section were determined by adding or subtracting the number of intervening sections (30 μm intervals). The analysis was performed as follows: 1) RTN/pFRG: 4 sections rostral from the caudal end of the facial nucleus (bregma: −6.48 to −6.12 mm); 2) NTS: 6 sections rostral from the caudal end of the most caudal NTS setion (bregma: −7.92 to −7.32 mm). For the RTN/pFRG level, the mapping was limited to the ventral half of the brainstem which contains the distinctive and isolated parafacial cluster of Phox2b-expressing neurons.

Sections were also examined to confirm that fiber optics were placed in the intended location. When fiber optics were not correctly positioned, these mice were excluded from the analysis.

### Statistics

Light-evoked changes in breathing were determined as follows. Respiratory frequency (electromyography or breathing flow) was measured during the minute preceding the photostimulation episode and was remeasured during the second minute after the end of the photostimulus, i.e., after full recovery. The average of these two values was considered as the baseline. The parameters were measured throughout the photostimulation episodes (5-20 Hz), and the effect of the light was calculated as a percentage increase from baseline. The effects of chemogenetic inhibition were examined in a similar fashion by comparing the values of the respiratory variables during the last 3 min of normoxia, hypercapnic or hypoxia exposure to the averaged pretest and posttest baseline. Results were compared using a Kruskal-Wallis One or Two Way ANOVA on ranks followed by all multiple comparisons (Bonferoni’s method). Differences were considered significant when p<0.05. In figures, data from each animal are presented in scatter plots, with summary data in text shown as mean ± SD.

## Results

### Histological characterization of Phox2b::cre;Ai32 mice

The histological experiments were designed to verify whether, in Phox2b::cre mice crossed with Ai32, there is selective expression of ChR2 in Phox2b-expressing neurons. In Phox2b::cre;Ai32 mice, 96.3 ± 2.1% of transduced (GFP-expressing) neurons had a Phox2b-immunoreactive nucleus within the RTN region (Figs. 1A-C), 89.7 ± 11.6% of transduced neurons had a Phox2b-immunoreactive nucleus in the carotid body (Figs. 3A-C), and 92.4 ± 5.2% of transduced neurons had a Phox2b-immunoreactive nucleus into the NTS (Fig. 4A). Within the ventrolateral medulla, the Phox2b-expressing neurons represent 26.1 ± 5.5% of TH-immunoreactive, i.e. catecholaminergic neurons (average for the five mice) (data not shown). In the carotid body, Phox2b-expressing neurons represent 97.8 ± 3.8% of TH-immunoreactive, while in the NTS region, they represent 16.1 ± 8.3% of TH-immunoreactive neurons.

### Photostimulation of Phox2b-expressing neurons in the RTN increases breathing frequency in anesthetized Phox2b::cre;Ai32 mice treated with fentanyl

We did not find significant differences between males and females in all experimental protocols; therefore, the data were combined.

In urethane-anesthetized Phox2b::cre;Ai32 mice, photostimulation (5, 10 and 20 Hz) of the RTN region produced a significant increase in breathing frequency (fR: 187.7 ± 17.7; 202.7 ± 20.04; 215.1 ± 17.5, vs. baseline: 166.3 ± 18.6 breaths/min; F_3,146_ = 236.5; p<0.0001) (Figs. 1D-L). The breathing stimulation increased as a function of the light pulse frequency (Figs. 1D-F) but did not vary according to the duration of the light pulses (1, 5 or 10 ms) (data not shown). Importantly, the effects of photostimulation on fR were very reproducible within each animal and we did not observe a significant run-down of the effect over the course of an experiment. Photostimulation (20 Hz) in control mice in which RTN neurons did not express ChR2 had no effect on breathing (165.8 ± 17.6, vs. baseline: 164.7 ± 17.9 breaths/min; p = 0.88). In the same preparation, ip injection of fentanyl (500 μg/kg) produced a roughly 53% decrease in baseline fR (Figs. 1D-I). Maximum breathing frequency suppression occurred 5-7 min following fentanyl administration. After 10 min, Phox2b-expresing neurons were selectively stimulated (5, 10 and 20 Hz) following fentanyl-induced reduction in fR. Optogenetic stimulation (5 and 10 Hz) of the Phox2b-expressing neurons in the RTN produced a significant increase in breathing frequency that reached the baseline levels before fentanyl-induced bradypnea but never reached the same level of RTN stimulation without fentanyl. For example, RTN photostimulation (5 and 10 Hz) increased fR (143.4 ± 6 and 159.1 ± 10.2, vs. baseline without fentanyl: 166.3 ± 18.6 breaths/min; F_2,68_ = 44.6; p = 0.036) (Figs. 1D-E, 1G-H). However, with the 20 Hz stimulation, the fR reached the same values of the photostimulation without fentanyl, i.e RTN stimulation at 20 Hz was able to have a further increase in breathing rate (213 ± 20.7, vs. 20 Hz without fentanyl: 215.1 ± 17.5 breaths/min; p = 0.93) (Figs. 1F and 1I). The change in fR produced by 10 Hz (54.7 ± 5.1, vs. saline: 43.9 ± 3.13 breaths/min; t = 9.63, df = 14, p = 0.0006) and 20 Hz (96.2 ± 8.6, vs. saline: 58.7 ± 5.8 breaths/min; t = 10.02, df = 14, p = 0.0006) photostimulation was greater under fentanyl-induced respiratory depression (Figs. 1K-L).

### Photostimulation of Phox2b-expressing RTN neurons increases breathing frequency in conscious Phox2b::cre;Ai32 mice treated with fentanyl

Photostimulation (5, 10 and 20 Hz) of Phox2b-expressing RTN neurons in conscious mice in room air (FiO_2_ = 0.21) produced a marked increase in fR (Figs 2A-I). Similar to anesthetized approach, the effects of optogenetic stimulation on fR were very reproducible within each animal over the course of the experiments. Photostimulation in control mice that do not express ChR2 in the RTN region had no effect on breathing (185.6 ± 11.2, vs. baseline: 187.1 ± 13.6 breaths/min; p = 0.13). Therefore, breathing stimulation produced by the optogenetic stimulation in the experimental group was dependent on the expression of ChR2 by the RTN neurons. The increase in fR was virtually instantaneous and lasted throughout the stimulus duration. The stimulation period was followed by a period of hypopnea (reduced ventilation) that varied in severity and duration according to the magnitude of the previous stimulus-evoked activation of fR (Figs. 2D-F).

**Figure 2:**
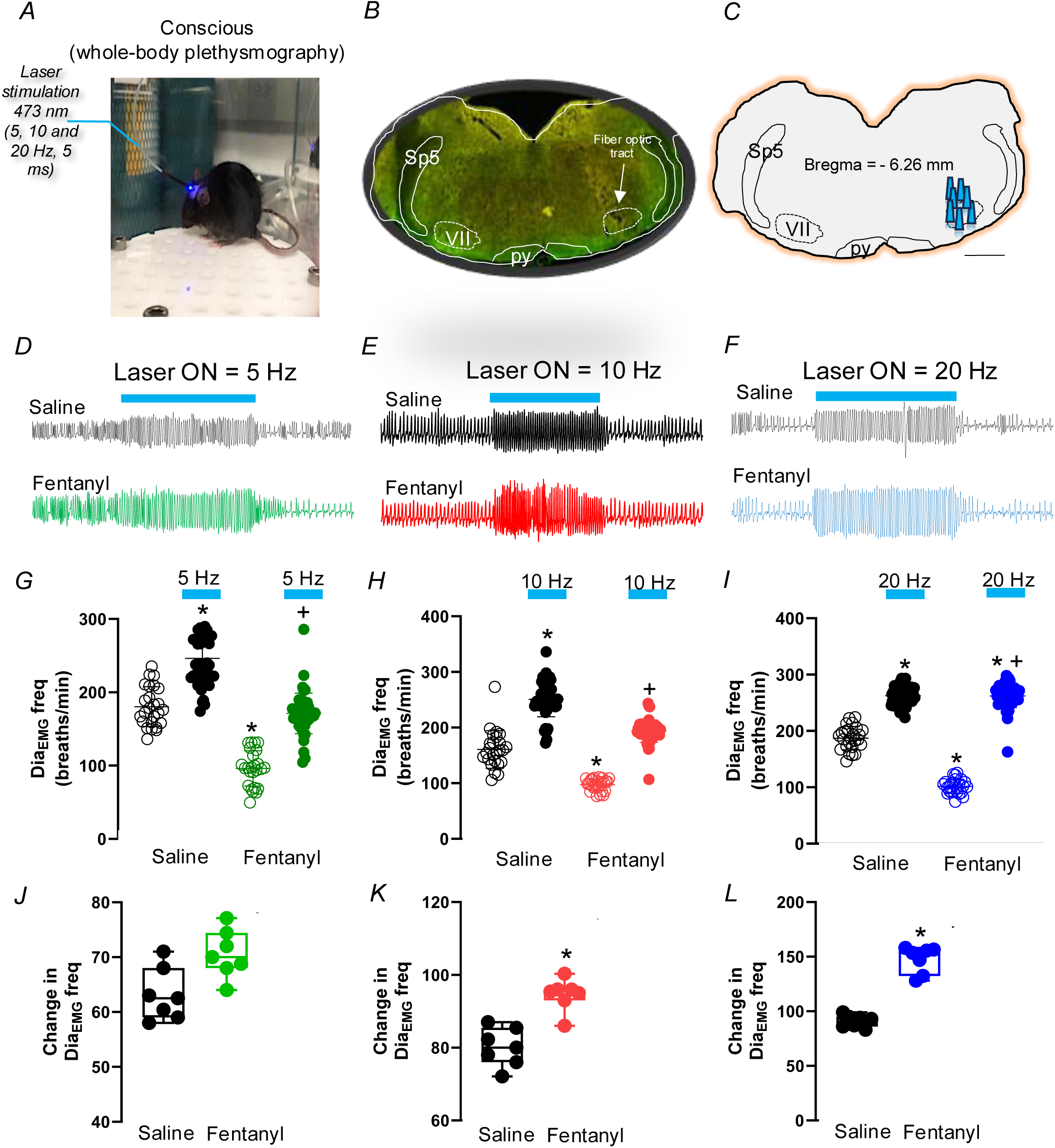
Optogenetic activation of ChR2-expressing Phox2b RTN neurons activates breathing in conscious mice treated with fentanyl. A) Schematic of the *in vivo* (conscious) preparations. In conscious mice, a previous optrode was implanted into the RTN and breathing activity was recorded through whole body plethysmography. B) Photomicrograph showing typical unilateral fiber optics tracts in the RTN region. C) Computer-generated plot of fiber optics placement that were confine to the RTN region (bregma −6.26 mm according to the Franklin and Paxinos Atlas) (41). D-F) Plethysmography recording of the respiratory effects of photostimulation (5, 10 and 20 Hz, 10 ms light pulse) of the Phox2b-expressing neurons in the RTN region after ip injection of saline or fentanyl (500 μg/kg). G-I) Group data for the respiratory frequency effect of G) 5Hz (One way repeated-measure ANOVA F_3, 146_ = 130.3; p<0.0001), H) 10 Hz (One way repeated-measure ANOVA F_3, 146_ = 209.2; p<0.0001), and I) 20 Hz photostimulation of Phox2b-expressing RTN neurons in a saline or fentanyl injection (One way repeated-measure ANOVA F_3, 146_ = 564.3; p<0.0001). J-L) Changes in respiratory frequency elicited by J) 5Hz, K) 10 Hz, and L) 20 Hz photostimulation of Phox2b-expressing RTN neurons in a saline or fentanyl injection. Abbreviations: py, pyramid tract; Sp5, spinal trigeminal nucleus; VII, facial motor nucleus. Scale bar = 1 mm, applied to B-C. *Different from baseline (saline); +Different from baseline (fentanyl).

Like anesthetized animals, the respiratory rate (fR) decreased by 55% as shown in Figures 2D-I. Optogenetic stimulation (5 and 10 Hz) of Phox2b-expressing neurons in the RTN significantly increased breathing frequency, returning it to baseline levels observed prior to fentanyl-induced bradypnea. However, these levels did not match those achieved by stimulating RTN without fentanyl. For instance, photostimulation at 5 and 10 Hz increased the fR to 171.3 ± 27.6 and 193.7 ± 20.13 breaths/min, respectively, compared to the baseline without fentanyl (246.3 ± 39.6 breaths/min; p = 0.0026), as shown in Figures 2D, 2G, 2E, and 2H. On the other hand, with 20 Hz stimulation, the fR reached levels comparable to those of photostimulation without fentanyl; that is, RTN stimulation at 20 Hz further increased the breathing rate to 262.1 ± 20.5, similar to 20 Hz stimulation without fentanyl (261.6 ± 16.3 breaths/min; p = 0.98), as indicated in Figures 2F and 2I. Moreover, the magnitude of increase in fR induced by 10 Hz and 20 Hz photostimulation was greater under conditions of fentanyl-induced respiratory depression (94.5 ± 4.3 vs. saline: 80 ± 5.2 breaths/min; t = 5.58, df = 14, p = 0.001 for 10 Hz and 147.3 ± 12.3 vs. saline: 90 ± 5.7 breaths/min; t = 11, df = 14, p<0.0001 for 20 Hz), as depicted in Figures 2K-L.

For stimulation experiments in conscious mice, a fiber optic (N = 7) was unilaterally implanted directly into the RTN region in Phox2b::cre;Ai32 mice. The location of the fiber optic was placed unilaterally in the RTN region (Figs. 2B-C). The injection center was 150-200 μm above the ventral medullary surface, 100 μm rostral to the caudal end of this nucleus, and 1.4 mm lateral to the midline, as previously demonstrated (43) (Figs. 2B-C).

### Optogenetic activation of ChR2-expressing Phox2b carotid body or NTS cells activates breathing frequency in mice treated with fentanyl

To optogenetically activate Phox2b-expressing cells in carotid body or NTS *in vivo*, we used the Phox2b::cre;Ai32 that encodes ChR2 in Phox2b neuronal profile (Figs. 3A and 4A)(36). Both the carotid body and the NTS express Phox2b neurons that are involved in respiratory regulation (44, 45). A robust activation of respiratory rate was evoked by photostimulation (5, 10 and 20 Hz) of the carotid body (247.5 ± 13, 263.6 ± 12.45 and 295.8 ± 9.7 vs. baseline: 158.6 ± 4.9 breaths/min; F_3,206_ = 250.1; p<0.0001) or the NTS (223.7 ± 30, 251.2 ± 36 and 264.2 ± 40.7 vs. baseline: 178.5 ± 33 breaths/min; F_3,146_ = 559.5; p<0.0001) (Figs. 3A-J and 4C-H).

**Figure 3:**
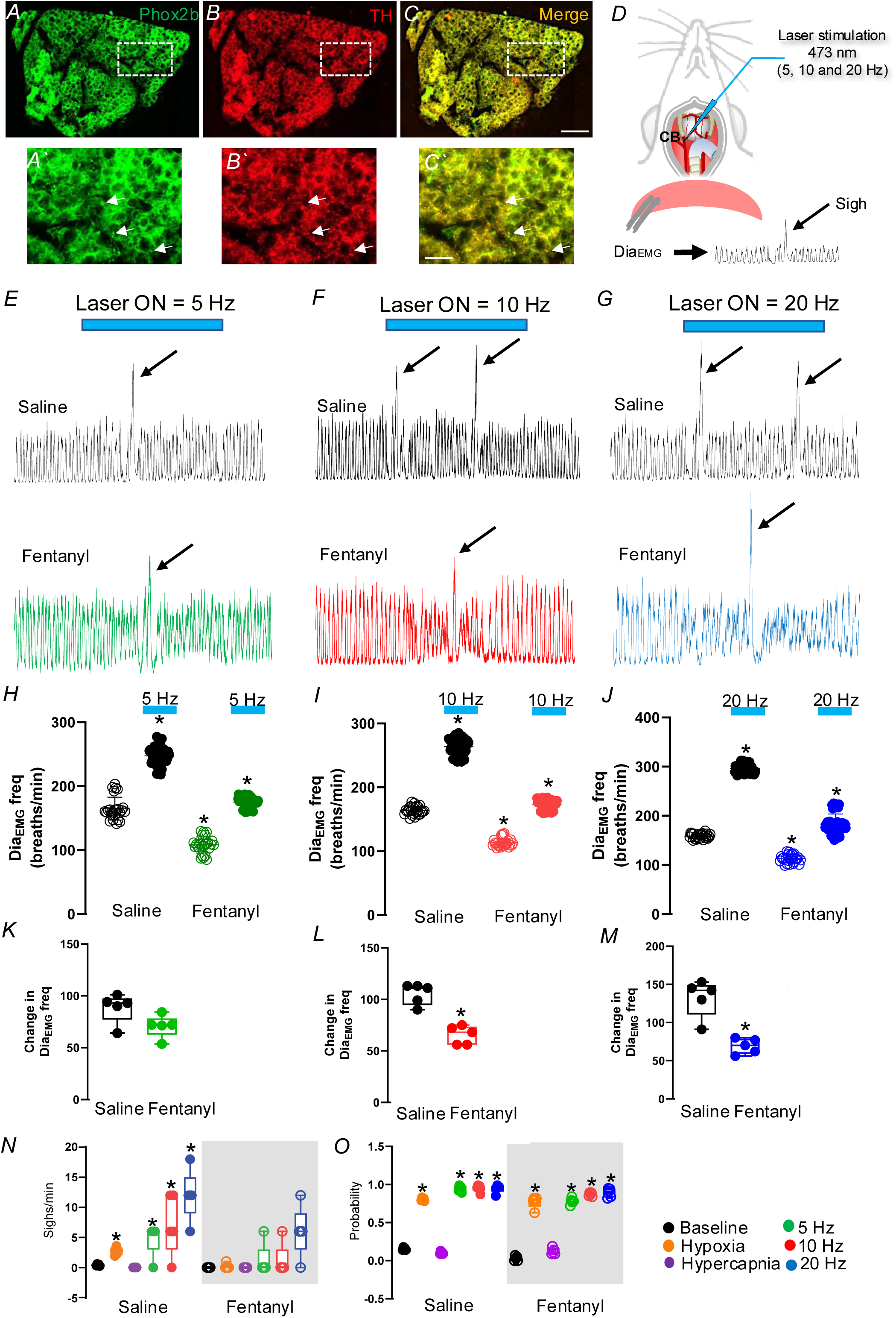
Optogenetic activation of ChR2-expressing Phox2b carotid body cells activates breathing in mice treated with fentanyl. A-C) Immunohistochemistry showing the co-localization of Phox2b fluorescence in TH-expressing cells of the carotid body. A’-C’) Higher magnification of the delineated square in A-C. D) Schematic of the *in vivo* (urethane anesthetized) preparation. In anesthetized mice, diaphragm muscle was recorded and a ventral optrode was placed in the carotid body. E-G) Representative recordings of diaphragm electromyography (Dia_EMG_) from an anesthetized mouse after ip injection of saline or fentanyl (500 μg/kg) under baseline or photostimulation stimulation (5, 10 and 20 Hz, 10 ms light pulse) of the Phox2b-expressing cells in the carotid body region. H-J) Group data for the Dia_EMG_ respiratory frequency effect of H) 5Hz (One way repeated-measure ANOVA F_3, 146_ = 772.3; p<0.0001), I) 10 Hz (One way repeated-measure ANOVA F_3, 146_ = 1782; p<0.0001), and J) 20 Hz photostimulation of Phox2b-expressing carotid body cells in a saline or fentanyl injection (One way repeated-measure ANOVA F_3, 146_ = 1267; p<0.0001). K-M) Changes in Dia_EMG_ respiratory frequency elicited by K) 5 Hz, L) 10 Hz, and M) 20 Hz photostimulation of Phox2b-expressing neurons of the carotid body region in a saline or fentanyl injection. N) Number of sighs generated under baseline, hypoxia (FiO_2_ = 0.1), hypercapnia (FiCO_2_ = 0.1) or photostimulation (5, 10 and 20 Hz, 10 ms light pulse) of the Phox2b-expressing neurons in the carotid body region after saline or fentanyl treatment (One way repeated-measure ANOVA F_11, 48_ = 137.22; p<0.0001). O) Probability of evoking a sigh by hypercapnia, hypoxia and optogenetic stimulation of Phox2b-expressing neurons of the carotid body (One way repeated-measure ANOVA F_11, 48_ = 383.3; p<0.0001). Scale bar = 200 μm, applied to A-C; 50 μm, applied to A’-C’. *Different from baseline (saline).

After fentanyl administration, photostimulation of carotid body (173.9 ± 7.6, 177.5 ± 7.2 and 183 ± 20.7) or the NTS (130.2 ± 29.6, 168.8 ± 28.7 and 176.1 ± 30.4) increased the fR compared to the baseline without fentanyl (163.7 ± 6.7 breaths/min) (Figs. 3E-M; 4C-K). Interesting to note is that the change in fR produced by 10 (65.4 ± 8.9, vs. saline: 105.2 ± 10.3 breaths/min; t = 6.53, df = 10, p = 0.002) and 20 Hz (69 ± 10, vs. saline: 132.2 ± 24.5 breaths/min; t = 5.34, df = 10, p = 0.0007) photostimulation of carotid body was smaller under fentanyl-induced respiratory depression (Figs. 3L-M). We did not find significant differences in fR if the NTS was stimulated before or after fentanyl (Figs. 4I-K).

**Figure 4:**
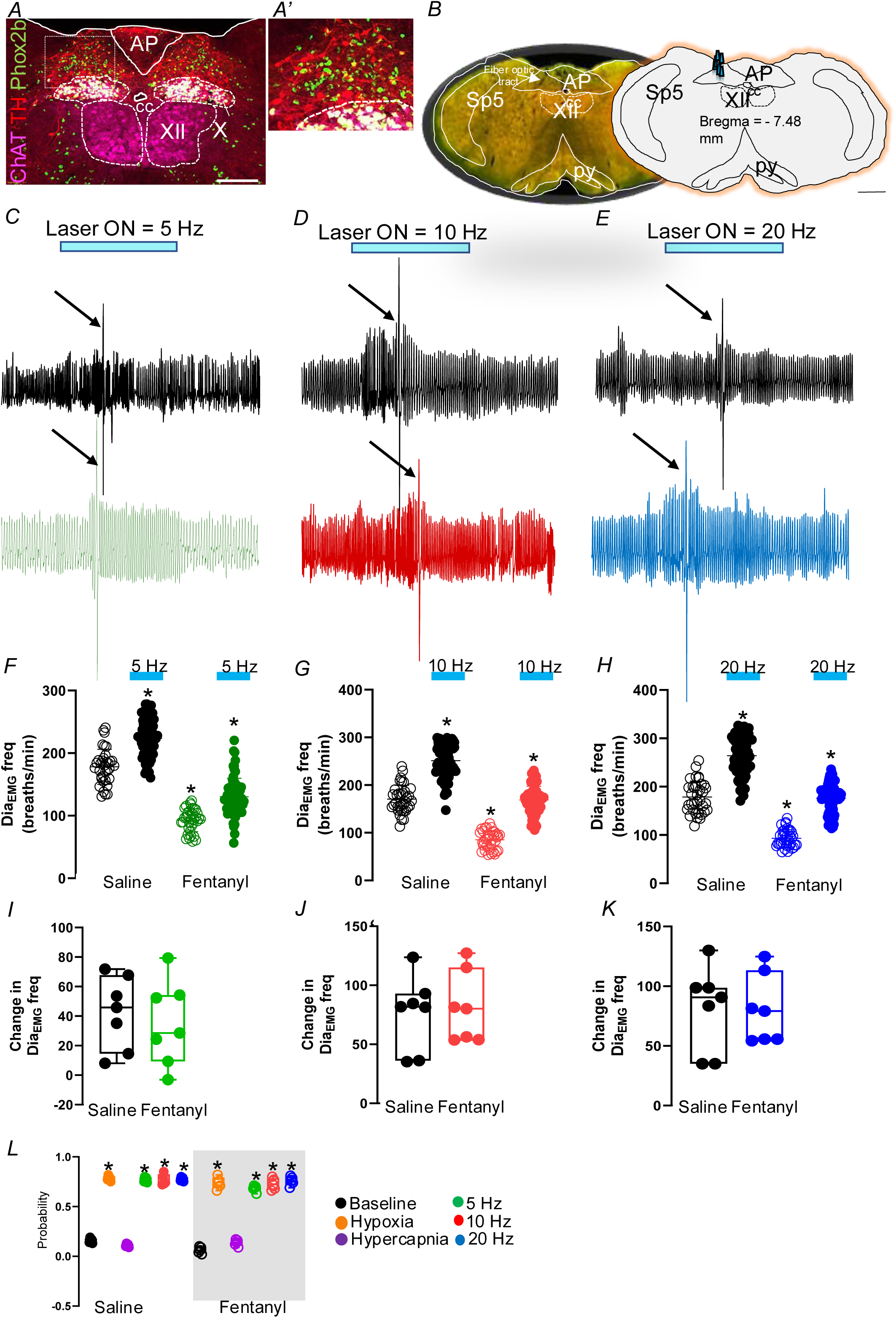
Optogenetic activation of ChR2-expressing Phox2b NTS neurons activates breathing in mice treated with fentanyl. A) Immunohistochemistry showing the co-localization of Phox2b fluorescence in the nucleus of the solitary tract. A’) Higher magnification of the delineated square in A. B) Photomicrograph showing typical unilateral fiber optics tracts in the NTS region. Computer-generated plot of fiber optics placement that were confined to the NTS region (bregma −7.48 mm according to the Franklin and Paxinos Atlas) (41). C-E) Plethysmography recording of the respiratory effects of photostimulation (5, 10 and 20 Hz, 10 ms light pulse) of the Phox2b-expressing neurons in the NTS region after ip injection of saline or fentanyl (500 μg/kg). F-H) Group data for the respiratory frequency effect of F) 5Hz (One way repeated-measure ANOVA F_3, 206_ = 218.4; p<0.0001), G) 10 Hz (One way repeated-measure ANOVA F_3, 206_ = 250.1; p<0.0001), and H) 20 Hz photostimulation of Phox2b-expressing NTS neurons in a saline or fentanyl injection (One way repeated-measure ANOVA F_3, 206_ = 221.8; p<0.0001). I-K) Changes in respiratory frequency elicited by I) 5 Hz, J) 10 Hz, and K) 20 Hz photostimulation of Phox2b-expressing neurons of the NTS in a saline or fentanyl injection. L) Probability of evoking a sigh by hypercapnia, hypoxia and optogenetic stimulation of Phox2b-expressing NTS neurons (One way repeated-measure ANOVA F_11, 48_ = 329.5; p<0.0001). Abbreviations: AP, area postrema; cc, central canal; py, pyramid tract; Sp5, spinal trigeminal nucleus; X, vagus nerve nucleus; XII, hypoglossal motor nucleus. Scale bar = 200 μm in A; 1 mm in B. *Different from baseline (saline).

### Increased sigh rate due to optogenetic activation of ChR2-expressing Phox2b carotid body or NTS cells are blunted after fentanyl

It is known that the sigh rate increases in hypoxia (17, 20, 46), and that both carotid body and NTS neurons are involved in the hypoxia pathway (16). To examine the response of sighing to carotid body or NTS stimulation, we optogenetically activated the Phox2b-expressing cells in the carotid body or in the NTS and compared with the sigh generated by hypoxia (FiO_2_ = 01) and hypercapnia (FiCO_2_ = 0.07). As expected, sighing increased immediately after the switch from normoxia to hypoxia conditions, from 0.42 ± 0.13 sighs/min in normoxia (FiO_2_ = 0.21), to 2.9 ± 0.24 sighs/min in hypoxia (FiO_2_ = 0.1) (Figs. 3E-G, 3N-O, 4C-E, and 4L). No sigh was observed after hypercapnia (0.002 ± 0.004, vs. normoxia: 0.42 ± 0.13 sighs/min; p>0.99) (Figs. 3J, and 4J). Interesting, fentanyl (500 μg/kg - ip) eliminated sigh activity elicited by hypoxia (0.003 ± 0.004, vs. hypoxia: 2.9 ± 0.24 sighs/min; p = 0.88), carotid body (6 ± 4.24, vs. Phox2b carotid body stimulation: 12 ± 4.24 sighs/min; p = 0.03) and NTS (1.2 ± 2.68, vs. Phox2b NTS stimulation: 6.8 ± 1 sighs/min; p = 0.032) neuronal stimulation, suggesting that sigh generation is impaired under opioid-induced respiratory depression (Figs. 3J, and 4J).

### Selective stimulation of *Nmb*-expressing RTN neurons increases breathing frequency in conscious mice treated with fentanyl

It is well described that in rodents, RTN neurons are distinguished by a unique combination of molecular markers, including the expression of Phox2b, vesicular glutamate transporter 2, tachykinin receptor 1, galanin, and, more recently, neuromedin B (*Nmb*) (33). Among these markers, *Nmb* has proven to be a reliable marker for identifying RTN neurons involved in respiratory regulation (43, 47). Having established the involvement of Phox2b-expressing neurons in the RTN during fentanyl-induced respiratory depression, our next series of experiments will delve deeper into the role of *Nmb*-expressing RTN neurons in mediating this respiratory depression in conscious mice.

Injection of two distinct AAVs encoding cre-dependent into the RTN of *Nmb*::cre^/+^ mice [AAV5-EF1a-hChR2(H134R)-mCherry (N = 8), and AAV5-hSyn-DIO-hM4D(Gi)-mCherry (N = 7)] resulted in labeling restricted to RTN *Nmb* neurons (97.5 ± 3.4% of labeled neurons contained *Nmb* mRNA), with transduction of 78 ± 4% of all VGlut2 neurons (Figs. 5A-C).

**Figure 5:**
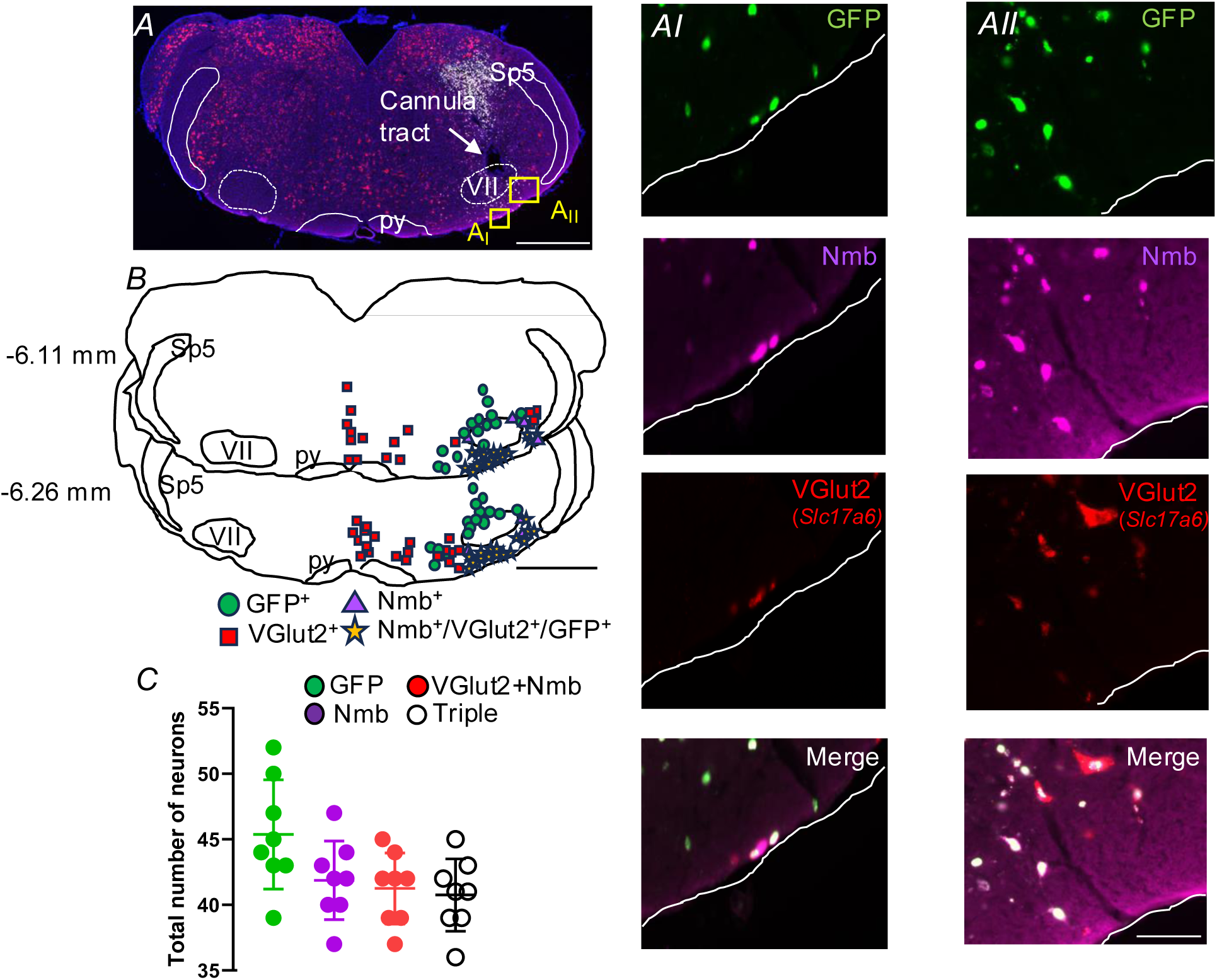
Selective expression of ChR2 in *Nmb*-expressing neurons of the RTN region. A) Photomicrography of combined *in situ* hybridization (ISH) (*Nmb* and VGlut2 neuronal profile) and immunohistochemistry (GFP) showing selective transduction of RTN neurons in a *Nmb*^::^cre mouse. AI and AII) Higher magnification of the yellow delineated square in A. B) Rostral-to-caudal series of transverse sections (bregma levels in mm as indicated) depicting the location of *Nmb*^+^ChR2-GFP-expressing neurons overlaid from 8 RTN-targeted cases. C) Total number of neurons within the RTN region Scale bar = 1 mm in A, applied A-B; 50 µm in AII, applied to AI-AII. Abbreviations: py, pyramid tract; Sp5, spinal trigeminal nucleus; VII, facial motor nucleus.

In conscious mice in room air (FiO_2_ = 0.21), photostimulation (5, 10 and 20 Hz) of *Nmb*-expressing RTN neurons produced a marked increase in f_R_ (Figs. 6A-F). Like the optogenetic stimulation of Phox2b-expressing neurons, the effects of stimulation on f_R_ were highly consistent within each animal throughout the experiments. Photostimulation in control mice that do not express ChR2 in the RTN region had no effect on breathing (184.6 ± 13, vs. baseline: 181.4 ± 19 breaths/min; p = 0.87). Optogenetic stimulation of *Nmb*-expressing neurons in the RTN increased f_R_ almost immediately and sustained this effect throughout the stimulation period (Figs. 6A-C). This was followed by a hypopnea phase, with severity and duration depending on the intensity of the prior stimulation. The response required ChR2 expression in the RTN neurons (Figs. 6A-C). As expected, fentanyl reduced the f_R_ by 50% and optogenetic stimulation (5 and 10 Hz) of *Nmb*-expressing neurons in the RTN significantly increased f_R_, returning it to baseline levels observed prior to fentanyl-induced reduction in respiratory rate (Figs. 6A-B and 6D-E). However, these levels did not match those achieved by stimulating RTN without fentanyl (Figs. 6D-E). For instance, photostimulation at 5 and 10 Hz increased the fR to 180 ± 36.7 and 172.2 ± 15.8 breaths/min, respectively, compared to the baseline without fentanyl (173.8 ± 29.1 breaths/min; p = 0.997) (Figs 6A-B and 6D-E). With twenty (20) Hz stimulation, the f_R_ reached levels comparable to those of photostimulation without fentanyl; for example, *Nmb*-neurons stimulation at 20 Hz further increased the breathing rate to 281.4 ± 30.4, similar to 20 Hz stimulation without fentanyl (282.6 ± 71,7 breaths/min; p = 0.99) (Figs. 6C and 6F). The magnitude of increase in f_R_ induced by 20 Hz photostimulation was greater under conditions of fentanyl-induced respiratory depression 208 ± 25.2 vs. saline: 129.9 ± 26 breaths/min; t = 6.794, df = 16, p<0.0001 for 20 Hz) (Figs. 6I).

**Figure 6:**
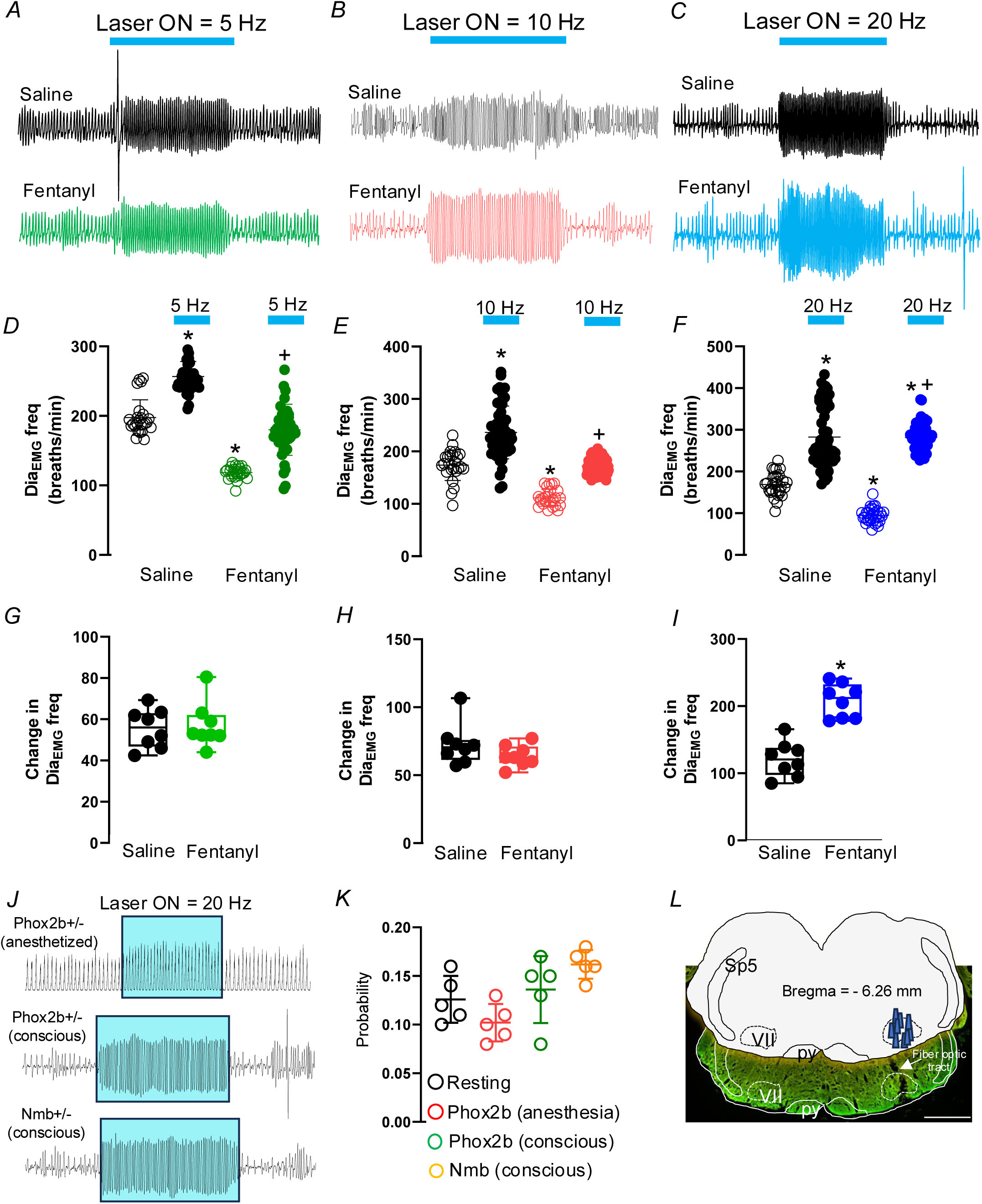
Optogenetic activation of ChR2-expressing *Nmb* RTN neurons activates breathing in conscious mice treated with fentanyl. A-C) Plethysmography recording of the respiratory effects of photostimulation (5, 10 and 20 Hz, 10 ms light pulse) of the *Nmb*-expressing neurons in the RTN region after ip injection of saline or fentanyl (500 μg/kg). D-F) Group data for the respiratory frequency effect of D) 5Hz (One way repeated-measure ANOVA F_3, 146_ = 158.8; p<0.0001), E) 10 Hz (One way repeated-measure ANOVA F_3, 146_ = 82.23; p<0.0001), and F) 20 Hz photostimulation of *Nmb*-expressing RTN neurons in a saline or fentanyl injection (One way repeated-measure ANOVA F_3, 176_ = 148.4; p<0.0001). G-I) Changes in respiratory frequency elicited by G) 5 Hz, H) 10 Hz, and I) 20 Hz photostimulation of *Nmb*-expressing RTN neurons in a saline or fentanyl injection. J) Dia_EMG_ or plethysmography recordings in Phox2b::cre mouse (urethane-anesthetized or conscious, respectively) and *Nmb*::cre mouse demonstrate the absence of sigh generation due to selective optogenetic stimulation of Phox2b- or *Nmb*-expressing cells in the RTN region. K) Probability of evoking a sigh by optogenetic stimulation of Phox2b- or *Nmb*-expressing RTN neurons (One way repeated-measure ANOVA F_3, 36_ = 1.18; p = 0.732). L) Photomicrograph showing typical unilateral fiber optics tracts in the RTN region. Computer-generated plot of fiber optics placement that were confined to the RTN region (bregma −6.26 mm according to the Paxinos and Franklin Atlas, 2012). Abbreviations: py, pyramid tract; Sp5, spinal trigeminal nucleus; VII, facial motor nucleus. Scale bar = 1 mm applied to L. *Different from baseline (saline); +Different from baseline (fentanyl).

For stimulation experiments in conscious mice, a fiber optic (N = 8) was implanted directly in the RTN region in *Nmb*::cre mice with previous injection of AAV5-EF1a-hChR2(H134R)-mCherry. The location of the fiber optic was placed unilaterally in the RTN region (Figs. 6L).

Stimulation of Phox2b- or *Nmb*-expressing neurons in the RTN did not induce sigh under normoxic conditions compared to the resting state, suggesting that RTN neurons are not involved in increased sigh frequency (Figs. 6J-K).

### Selective inhibition of RTN *Nmb*-expressing neurons on breathing in conscious mice treated with fentanyl

We next used the chemogenetic strategy for genetically targeted inhibition of the RTN *Nmb* neurons with AAV5-hSyn-DIO-hM4D(Gi)-mCherry (Fig. 7A). The transduction rate was consistent with that shown in Figure 5C, and histological analysis indicates that hM4D(Gi)-mCherry-mediated transduction of RTN *Nmb* neurons in *Nmb*::cre^/+^ mice is both efficient and highly selective (Fig. 7B). Functional experiment showed a significant reduction in fR in RTN *Nmb*-inhibition with CNO (1 mg/kg, 0.2 ml) compared with saline (89.75 ± 4.3, vs. saline: 193.9 ± 6.38 breaths/min; p<0.0001) (Figs. 7C-D). Under the condition of inhibition of RTN *Nmb* neurons, the injection of fentanyl (500 μg/kg, ip) caused a significant further decrease (67.3 ± 3.95, vs. CNO: 89.75 ± 4.3 breaths/min, p = 0.034) in breathing frequency (Figs. 7C-D). A detailed examination of the breathing pattern revealed a distinctive phenotype in RTN *Nmb*-inhibited mice treated with fentanyl (Fig. 7C-D and 7H-I), including a marked increase in respiratory cycle duration variability and a higher incidence of apneas (10.07 ± 2.78, vs. control: 1.89 ± 0.32 apneas/hour, t = 8.26, df = 14, p<0.0001) (Figs. 7C and 7H-J). This was evident when comparing the respiratory traces, which showed a highly regular breathing pattern in control mice versus a more erratic pattern in RTN *Nmb*-inhibited mice treated with fentanyl, along with a greater dispersion of data points in the Poincaré plots (Figs. 7C and 7H-I).

**Figure 7:**
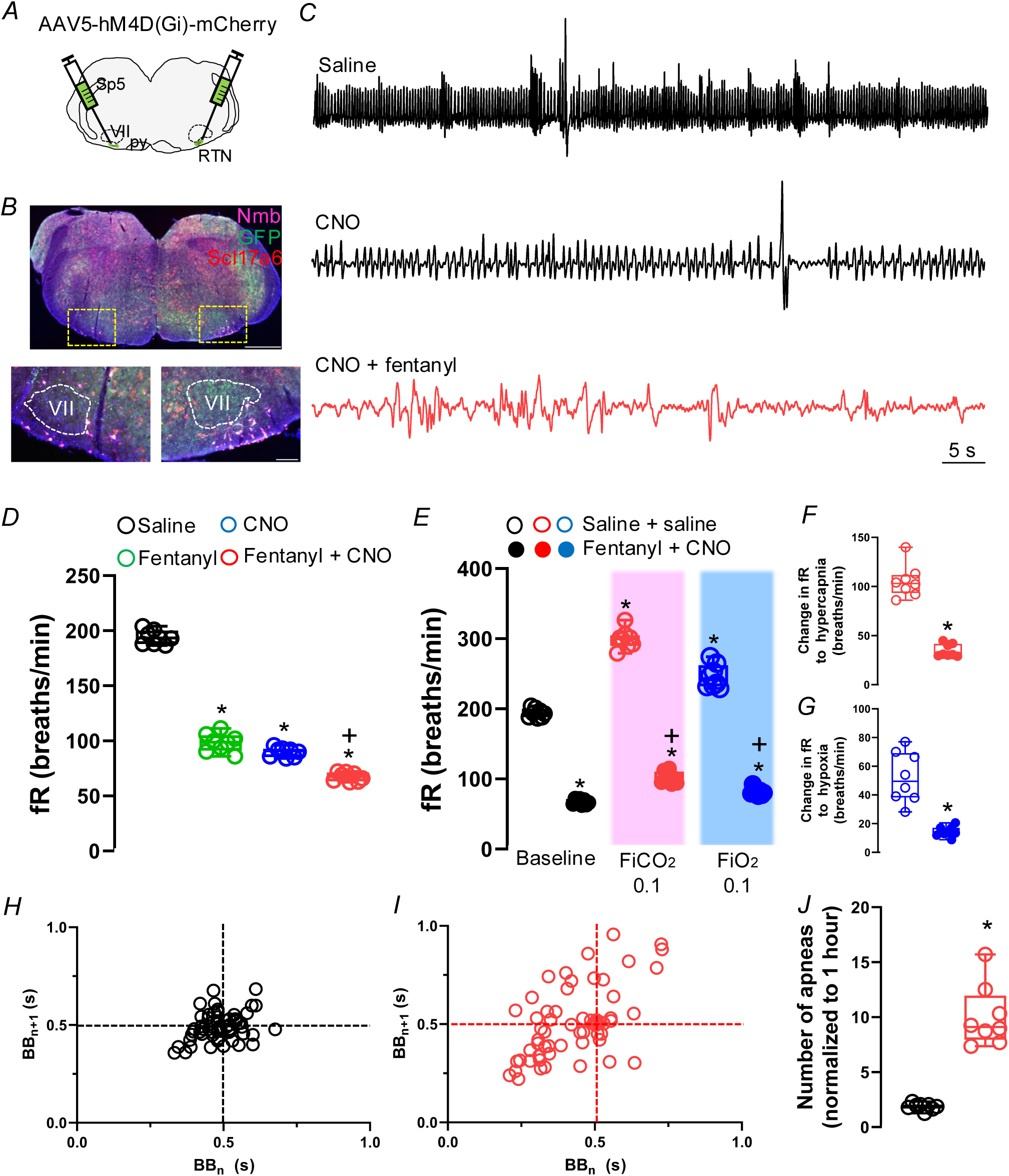
Selective inhibition of *Nmb*-expressing RTN neurons impairs breathing in conscious mice treated with fentanyl. A) Experimental strategy for genetically targeted inhibition of RTN *Nmb* neurons with AAV5-hM4D(Gi)-mCherry. B) Transverse section of mouse medulla showing *Nmb*-expressing RTN neurons labeled with a Cre-dependent mCherry. Viral transduction was confined to the RTN region. C) Plethysmography recording of the respiratory effects of saline (control) or CNO injection (1 mg/kg, ip) of the *Nmb*-expressing neurons in the RTN region after ip injection of saline or fentanyl (500 μg/kg). D) Group data for the respiratory frequency effect of saline (control) or CNO injection (1 mg/kg, ip) on *Nmb*-expressing neurons in the RTN region after injection of saline or fentanyl (500 μg/kg) (One way repeated-measure ANOVA F_3, 28_ = 696.8; p<0.0001). E) Group data for the respiratory frequency effect at normoxia, hypercapnia (FiCO_2_ = 0.1), or hypoxia (FiO_2_ = 0.1) on RTN *Nmb*-inhibiting neurons after injection of saline or fentanyl (500 μg/kg) (Two way repeated-measure ANOVA F_5, 35_ = 737.6; p<0.0001). F) Changes in respiratory frequency induced by hypercapnia (FiCO_2_ = 0.1) following RTN *Nmb* inhibition, after the administration of saline or fentanyl (500 μg/kg) (unpaired t-test = 11.34, df = 14; p<0.0001). G) Changes in respiratory frequency induced by hypoxia (FiCO_2_ = 0.1) following RTN *Nmb* inhibition, after the administration of saline or fentanyl (500 μg/kg) (unpaired t-test = 5.979, df = 14; p<0.0001). H-I) Representative Poincaré plots showing the breath duration (BBn) in relation to the subsequent breath duration (BBn+1) in control (black) (H) and RTN *Nmb*-inhibition under fentanyl (red) (I) mice. J) Number of apneas in control and RTN *Nmb*-inhibition under fentanyl mice (unpaired t-test = 8.266, df = 14; p<0.0001). Abbreviations: py, pyramid tract; Sp5, spinal trigeminal nucleus; VII, facial motor nucleus. *Different from baseline (saline); +Different from baseline (fentanyl).

Interestingly, both the tachypneic response to hypercapnia (101.9 ± 8.43, vs. control: 299.2 ± 13.9 breaths/min, t = 11.34, df = 14, p<0.0001) and hypoxia (81.7 ± 5.4, vs. control: 246 ± 16.7 breaths/min, t = 5.979, df = 14, p<0.0001) were significantly blunted in RTN *Nmb*-inhibited mice under fentanyl compared to controls, indicating that the RTN is essential for maintaining respiratory activity, particularly under OIRD (Figs. 7E-G).

## Discussion

In this study, we provide evidence of the respiratory effects mediated by selective activation of Phox2b-expressing neurons of the CB, NTS, and Phox2b or *Nmb-*expressing RTN neurons in the context of opioid-induced respiratory depression (OIRD). Our findings demonstrate that targeted stimulation of RTN neurons can significantly ameliorate the respiratory suppression caused by fentanyl, while stimulating CB or the NTS can elevate, but not ameliorate the fentanyl-induced respiratory depression. In addition, under fentanyl-induced respiratory depression, inhibition of the RTN results in irregular breathing patterns and a significant increase in the number of apneas, which disrupts respiratory homeostasis. This irregularity in breathing, coupled with frequent apneas, indicates that RTN activity is essential for stabilizing respiratory activities, especially under the suppressive effects of opioids. The compromised respiratory pattern highlights the RTN critical role in maintaining consistent ventilation, as its inhibition under fentanyl severely undermines the body’s ability to regulate breathing effectively.

Our data not only advances our understanding of the neural circuits involved in respiratory control but also opens new therapeutic avenues for potentially overcoming the OIRD specifically through stimulation of RTN neurons.

### ChR2-expressing neurons were likely RTN cells

The expression of channelrhodopsin in Phox2b or *Nmb*-expressing neurons provides significant insights into the functional architecture and potential role of these neurons, which are likely part of the RTN. The RTN has been implicated as a crucial site for chemosensation and the regulation of breathing (31, 32, 48), and our findings contribute to a deeper understanding of its neuronal components and their modulatory effects on respiratory rhythm.

The expression of Phox2b, a transcription factor, is a defining characteristic of neurons within the RTN. Prior studies have established Phox2b as a critical marker for the development and functional specification of the RTN (26, 27, 33, 35, 49–51). Our use of channelrhodopsin to manipulate the activity of these neurons allowed us to observe changes in respiratory patterns in response to photostimulation, affirming the role of these neurons in respiratory control. Specifically, the photostimulation led to an increase in breathing frequency, which correlates with the hypothesized role of RTN neurons in detecting and responding to CO_2_ concentrations (26, 29, 52). This observation supports the view that Phox2b-expressing neurons in the RTN play a pivotal role in the chemoreflex control of breathing. The expression of Phox2b coincided with other known markers of RTN neurons, such as NK1 receptors, the expression of *Nmb* on glutamatergic neurons, as well as the proton-activated G-protein coupled receptor GPR4 and the proton-inactivated potassium channel TASK-2, which have been shown to be involved in the neural circuitry controlling respiratory reflexes (52–54).

### RTN neurons rescue breathing under fentanyl-induced respiratory depression

The findings from our study demonstrate that selective stimulation of RTN neurons (Phox2b or *Nmb*-expressing cells) can rescue breathing patterns in the context of fentanyl-induced respiratory depression. This has significant implications for our understanding of respiratory control and potential therapeutic interventions for OIRD.

Our data clearly show that targeted activation of RTN neurons effectively counteracts the respiratory suppression typically observed with fentanyl administration. Previous research has identified the RTN as a critical hub for sensing CO_2_ levels and modulating breathing (23–25, 31, 32, 55). The ability of RTN neurons to override opioid-induced inhibition likely stems from their central role in maintaining homeostatic respiratory drive and the fact that these neurons do not express μ-opioid receptors (24, 30). When these neurons were activated in our experiments, there was a marked increase in both the rate and regularity of breathing, even in the presence of high-dose fentanyl, which typically depresses central respiratory drive (9, 56–58).

The reversal of fentanyl-induced respiratory depression by RTN neurons suggests an inherent capability of these neurons to bypass or override opioid-mediated inhibition pathways (59). Opioids primarily exert their respiratory depressing effects by binding to μ-opioid receptors in the brainstem, leading to a reduction in the responsiveness of central respiratory chemoreflexes. RTN neurons, known for their intrinsic chemosensitive properties, seem capable of overcoming this inhibition, possibly through an indirect or direct reactivation of the suppressed neural pathways involved in respiratory drive.

The ability of RTN neurons to restore respiratory function in opioid-compromised states holds promising clinical implications. Respiratory depression remains the most dangerous side effect of opioid analgesia or recreation use and is a leading cause of death in opioid overdoses. Our data suggest that opioids suppress the hypercapnic response presynaptic to the RTN which is likely an important factor in the events leading to the OIRD and the fatal consequences. The fact that the missing chemosensitive drive can be overcome by the modulation of RTN neuron activity presents a potential therapeutic pathway to mitigate risks associated with opioid therapy, especially in a clinical setting. Developing pharmacological or neuromodulation therapies targeting these neurons could offer a method to prevent or reverse OIRD without affecting the analgesic benefits of opioids.

It is also important to point out that 50% of preBötzinger neurons (key region for OIRD) do not express opioid receptors, which highlights the heterogeneous nature of this neuronal population (60, 61). This heterogeneity could mean that a subset of these neurons remains functional even during opioid exposure, which might partially explain the variability in individual responses to OIRD.

While the results are promising, our study has limitations that need to be addressed in future research. First, the specificity and efficiency of RTN neuron stimulation need further optimization. Our approach, while effective, might activate neighboring neurons or pathways, potentially confounding the exact mechanisms by which respiratory activity is restored. Furthermore, the long-term effects and safety of such interventions have yet to be thoroughly evaluated.

Future studies should focus on refining techniques for specific activation of RTN neurons and understanding the interaction between these neurons and opioid receptors. Additionally, exploring the genetic markers and molecular drivers of RTN neuron activity could provide deeper insights into their role in respiratory control and their potential modulation by various substances, including opioids.

### Role of Neuromedin B-expressing RTN neurons in respiratory regulation

Neuromedin B-expressing neurons in the RTN play a critical role in maintaining respiratory homeostasis (43). Ablation of these neurons has been shown to cause alveolar hypoventilation and, under fentanyl-induced respiratory depression, to result in irregular breathing patterns with increased variability (43), and present results. This suggests that *Nmb* neurons in the RTN are essential for stabilizing breathing under both normal and compromised conditions.

Periodic breathing, characterized by oscillations between hyperventilation and apnea, is a hallmark of respiratory instability observed in both humans and experimental models. This phenomenon is influenced by interactions among the apneic threshold, the sensitivity of the hypercapnic ventilatory response (HCVR, also referred to as loop gain), transient arousals, and cerebral blood flow regulation (62, 63). In the context of RTN *Nmb*-inhibited mice, an elevated apneic threshold is a particularly plausible contributor to breathing instability. These mice are unresponsive to CO_2_, aligning with earlier findings that RTN lesions in rats raise the apneic threshold (27).

Enhanced peripheral chemoreceptor activity and sensitivity further contribute to breathing instability in RTN *Nmb*-lesioned mice, mirroring mechanisms of periodic breathing observed in humans and dogs (63). The carotid body, which detects changes in O_2_, CO_2_, and H^+^ (64), is likely the primary source of compensatory respiratory drive in these mice, although contributions from aortic bodies and brainstem mechanisms cannot be excluded (65–67).

The instability in breathing observed in RTN *Nmb*-lesioned mice may therefore result from an interplay of factors: an elevated apneic threshold, heightened carotid body activity, transient arousal, and possibly altered cerebral blood flow secondary to respiratory dysfunction. This demonstrates the RTN critical role in integrating central and peripheral respiratory inputs to maintain stable breathing patterns.

### Carotid body and NTS neurons did not rescue breathing under fentanyl-induced respiratory depression

Despite the critical roles of both the carotid body and Phox2b-expressing neurons in the NTS in the hypoxic response and in the normal respiratory and cardiovascular regulation, and in contrast to the RTN, our findings suggest that neither the carotid body nor the Phox2b neurons in the NTS can rescue breathing during OIRD.

The carotid bodies contain the peripheral chemoreceptors that primarily respond to changes in blood oxygen, carbon dioxide, and pH levels, stimulating reflexive adjustments in respiratory activity (12, 13, 16). While they are pivotal in the body’s immediate response to hypoxemia, their role in response to hypercapnia and the modulation of respiratory patterns under opioid influence appears to be limited. Previous studies have shown that opioids can blunt the ventilatory response to CO_2_ by directly affecting central chemoreceptive sites rather than peripheral sensors like the carotid bodies (68). Our findings align with these observations, suggest that the carotid bodies, per se, may not be sufficient to overcome the central respiratory depression caused by fentanyl. This may possibly be due to a lack of influence over central opioid pathways that are heavily suppressed during opioid toxicity, or direct inhibition from opioid exposure (68, 69).

Phox2b-expressing neurons within the NTS play a critical role in integrating sensory input from peripheral chemoreceptors and baroreceptors and modulating autonomic functions accordingly (21, 70, 71). However, under conditions of fentanyl-induced depression the integration and processing functions of the NTS appear to be compromised. This could be due to the potent inhibitory effects of opioids on neurotransmitter release and neuronal excitability within the central brainstem circuits, which include the NTS (71).

The inability of the carotid body and Phox2b neurons in the NTS to counteract OIRD poses a critical challenge in addressing this condition. This necessitates further exploration into alternative strategies that could potentially bypass or counteract the suppressive effects of opioids on central respiratory pathways. The RTN, as discussed in earlier sections, offers a promising target given its apparent capacity to override opioid suppression through direct chemoreception and neural activation.

Future studies should aim to elucidate the specific mechanisms through which fentanyl and other opioids inhibit the carotid body and NTS functions. Additionally, investigating the potential compensatory mechanisms within other parts of the respiratory network, such as the pons or the pre-Bötzinger complex, could provide further insights into strategies for managing OIRD (56, 58, 72, 73). Genetic and pharmacological manipulations, alongside advanced imaging techniques, could be pivotal in uncovering these mechanisms.

### Sigh Generation: The Role of Phox2b-Expressing Cells in the Carotid Body and Nucleus of the Solitary Tract, and *Nmb* Neurons in the Retrotrapezoid Nucleus

Optogenetic stimulation of defined neural circuits has advanced our understanding of the physiological and pathological mechanisms governing respiratory control, including the regulation of sigh generation. The physiological sigh has been suggested to play a critical role in the hypoxic response and in preventing alveolar collapse and contributing to central respiratory drive (46). Thus, inhibition of sigh activity from opioid exposure represents an enhanced state of vulnerability to respiratory pathology. The carotid body and the NTS are two critical components in the respiratory network, influencing breathing patterns in response to various internal and external stimuli (18, 21, 74, 75). Recent experiments targeting both structures have deepened our understanding of the neural control of sighing, particularly under conditions of hypoxia (18).

We showed that optogenetic activation of Phox2b-expressing cells in the carotid body or the NTS increases the frequency of sighs. This effect aligns well with the known response of these cells to hypoxic conditions, wherein an increase in sigh rate helps to re-inflate alveoli and enhance oxygenation. This mechanism suggests that the respiratory network is wired to naturally enhance sigh production through these pathways when confronted with reduced oxygen levels.

The carotid body acts as a peripheral chemoreceptor, sensing changes in blood oxygen levels and communicating this information to the brainstem, which adjusts breathing patterns accordingly (12, 74, 76). Similarly, the NTS, which receives sensory input from the carotid body, plays a pivotal role in integrating and processing respiratory and cardiovascular signals (21, 22). Thus, optogenetically driving Phox2b-expressing cells in these areas artificially mimics the response to hypoxia, validating the essential role of these circuits in the control of sighing.

Interestingly, the sigh response induced by optogenetic activation of these pathways is significantly blunted under the influence of fentanyl. It is likely that fentanyl interferes with the neural transmission and responsiveness of the respiratory network, including the Phox2b-expressing cells in the carotid body and NTS. This blunting effect can be attributed to the action of fentanyl on opioid receptors, which are widely distributed in the central nervous system, including the NTS and carotid body (77, 78). By activating these receptors, fentanyl likely decreases the excitability of neurons or astrocytes responsible for initiating sighs, thereby overriding the stimulatory effects of optogenetic activation. This OIRD and loss of sigh represents a significant clinical challenge, especially in settings involving the risk of hypoxaemia, increasing the risk of breathing collapse.

RTN lesions increase sigh frequency at rest in both rats and mice, as a consequence of hypoxemia, since hyperoxia restores sigh frequency to control levels in mice (43). These findings suggest that RTN neurons may not play a direct role in hypoxia-induced sighs. However, other evidence indicates that a subset of RTN *Nmb*-expressing neurons regulates emotional sighing (20). Together, these data imply that after RTN neuron lesions, hypoxemia activates oxygen-sensitive mechanisms that drive breathing and partially compensate for reduced alveolar ventilation. While *Nmb* release from RTN neurons has been proposed to contribute to emotional sighing (17, 20), direct evidence using downregulation of *Nmb* in RTN neurons is still lacking. Moreover, our data show that selective activation of *Nmb*-expressing neurons was insufficient to induce sighs.

### Clinical implications

In our study, the dose of fentanyl administered induced approximately 50% reduction in breathing activity. This level of respiratory depression was chosen because it represents a clinically relevant scenario, providing a realistic framework for understanding the potential impacts of opioid administration in a medical setting and the efficacy of interventions aimed at mitigating OIRD. A 50% reduction in ventilation due to opioid administration is significant, as it mirrors situations commonly encountered in clinical environments. In such settings, opioids, particularly fentanyl, are frequently used for their potent analgesic properties. However, the narrow therapeutic window of opioids can lead to complications, primarily respiratory depression, which is a leading cause of morbidity and mortality associated with opioid overdose. The selected dose of fentanyl that results in a 50% decrease in ventilation effectively simulates moderate to severe respiratory depression while avoiding the immediate life-threatening conditions of complete respiratory failure, which is difficult to obtain within rodent and other non-human mammals. Moreover, the degree of respiratory depression elicited within this study allows for the examination of both the direct effects of opioids on the respiratory system and the efficacy of various interventions without the confounding factors introduced by total respiratory collapse. Understanding the effects of this specific level of respiratory suppression is crucial for developing targeted interventions. It provides a benchmark against which the effectiveness of treatments can be measured. For example, the most appropriate potential application of this optogenetic methodology could rescue breathing and the activation of airway dilator muscles, improving upper airway patency. Therefore, if an intervention can significantly restore ventilation in this model, it suggests potential utility in clinical settings, where reversing even partial opioid-induced respiratory depression can substantially improve patient outcomes. Behavioral tests designed to ascertain that such stimulation does not cause any untoward sensations or side effects would also be required.

### Conclusion

In conclusion, our study provides evidence that selective activation of Phox2b or *Nmb*-expressing neurons in the RTN, but not the CB or the NTS can significantly counteract fentanyl-induced respiratory depression. Additionally, the ability for RTN stimulation to enhance ventilation may even be enhanced during OIRD as evidenced by a greater magnitude of change in ventilation during RTN stimulation following fentanyl administration. This discovery advances our understanding of the neural circuits involved in respiratory control and highlights the RTN’s role in respiratory drive under opioid influence. By demonstrating that RTN neurons can restore breathing patterns even in the presence of high-dose fentanyl, we open new therapeutic avenues for managing opioid-induced respiratory depression, a major cause of mortality associated with opioid overdose.

## Acknowledgements

Supported by public funding from São Paulo Research Foundation (FAPESP) (Grants: 2022/07705-9 to ACT; 2022/09734-6 to TSM), Conselho Nacional de Desenvolvimento Científico e Tecnológico (CNPq) fellowship (306580/2023-3 to ACT, and 306418/2023-1 to TSM). This work was also funded by grants from the National Institutes of Health, National Heart, Lung, and Blood Institute: P01 HL14454 Project 2, R01HL151389, R01HL126523, R01HL144801 to JMR, and K99HL168211 to NJB.

